# Mechanisms for multiple activity modes of VTA dopamine neurons

**DOI:** 10.1101/008920

**Authors:** Andrew M. Oster, Philippe Faure, Boris S. Gutkin

## Abstract

Midbrain ventral segmental area (VTA) dopaminergic neurons send numerous projections to cortical and sub-cortical areas, and diffusely release dopamine (DA) to their targets. DA neurons display a range of activity modes that vary in frequency and degree of burst firing. Importantly, DA neuronal bursting is associated with a significantly greater degree of DA release than an equivalent tonic activity pattern. Here, we introduce a single compartmental, conductance-based computational model for DA cell activity that captures the behavior of DA neuronal dynamics and examine the multiple factors that underlie DA firing modes: the strength of the SK conductance, the amount of drive, and GABA inhibition. Our results suggest that neurons with low SK conductance fire in a fast firing mode, are correlated with burst firing, and require higher levels of applied current before undergoing depolarization block. We go on to consider the role of GABAergic inhibition on an ensemble of dynamical classes of DA neurons and find that strong GABA inhibition suppresses burst firing. Our studies suggest differences in the distribution of the SK conductance and GABA inhibition levels may indicate subclasses of DA neurons within the VTA. We further identify, that by considering alternate potassium dynamics, the dynamics display burst patterns that terminate via depolarization block, akin to those observed in vivo in VTA DA neurons and in substantia nigra pars compacta DA cell preparations under apamin application. In addition, we consider the generation of transient burst firing events that are NMDA-initiated or elicited by a sudden decrease of GABA inhibition, that is, disinhibition.

## Introduction

Dopamine is a key neurotransmitter involved in motivation, memory, and motor function. Two areas in the midbrain are particularly associated with large populations of dopaminergic neurons whose activity transmits dopamine signals: the ventral tegmental area (VTA) and the substantia nigra pars compacta (SNc). The dopamine signal originating from the SNc DA neurons is a necessary component for motor initiation [1–4], and the VTA DA signal is conjectured to convey reward-related information [5, 6].

The DAergic neurons display a wealth of dynamic behavior: pacemaker-like firing (seen in slice preparations), periods of burst activity, quiescence (due to either strong hyperpolarization or depolarization), and also sporadic activity (see e.g. [7–9]). In this paper, we examine the mechanisms underlying the range of activity modes found in DA neurons. Given the rich behavior of DA neuronal dynamics and due to their biological importance, a number of modeling efforts have been made to understand DA neural dynamics. Existing models can capture a variety of these behaviors [10–12], yet are not without their shortcomings. The models of [10–12] rely on somatodendritic interactions as underlying burst generation, however, two laboratories have experimental findings that suggest burst firing is somatically induced [13, 14]. These somatically induced bursts have characteristics consistent with normal bursting, suggesting that a single-compartmental model should be sufficient for generating the observed DA neuronal dynamics.

Here we present a single compartment model of a VTA DA neuron, which captures the essential qualitative behavior of DAergic neuronal dynamics. This revised model improves our earlier work [15] and better captures DA neuronal dynamics. Other single compartment models have been recently introduced [15–17], and these models differ from the presented model. For instance, the dynamics of the Ha-Kuznetsov model [17] predict a sustained fast firing regime under NMDA-bath-like conditions, which is not inline with the endogenous bursting patterns observed under NMDA-bath application [18]. In this work, we consider a range of conditions not considered in [15, 16], and we uncover dynamics that were heretofore undiscovered by modeling efforts. For instance, by considering alternate potassium dynamics, we observe burst patterns that terminate via depolarization block. Transient firing patterns have been observed in vivo in VTA DA neurons [19–21] and endogenous bursting of this form has been found in SNc DA cell preparations with apamin applied [22]. By performing numerical simulations for a variety of parameter sets, our model leads to a number of experimental predictions. Our studies suggest that neurons with a relatively low SK conductance fire in a high firing mode, are correlated with burst firing, and require higher levels of applied current before undergoing depolarization block. We also consider the role of GABAergic inhibition on an ensemble of neural types and examine alternate scenarios for both NMDA-initiated and disinhibition-initiated burst firing. Moreover, we may have unveiled another class of DA neurons that enters very fast firing regimes (though such a parameter regime may prove to be unphysiological).

The generation of DA neuronal activity patterns is dependent on a wide number of factors. At the most basic level, the DA pattern is dependent on both the intrinsic properties of the cells and the network activity that innervates the DA cells. Mameli-Engvall et al. recorded and observed an ensemble of activity modes that mouse VTA DAergic neurons express *in vivo* [23], shown in Fig. 1. Mameli-Engvall et al. classified neurons by their firing patters as either low or high firing (displaying a frequency of less than or greater than 5Hz, respectively) and further categorized neurons as either low or high bursting modes (less than 20% being low burst firing and high bursting being greater than 20% of spikes within burst events). Differences in the activity modes are evident when plotting the distributions of the associated interspike intervals, shown in Fig. 1b. In addition, [19] presents similar histograms for examining VTA DA neural ensembles. In this work, we examine the mechanisms behind this ensemble of dynamical classes of DA neurons. In Fig. 1c, we present the firing modes from our computational model for a neural ensemble that receives a wide-ranging amount of drive and containing neurons with weak to strong SK conductance. The model results expose a high correlation of weak SK conductance with the high firing regime (both bursting and tonic). Moreover, neurons with weak to medium SK conductances are also associated with a significant portion of the burst firing events. In fact, it is the balance of L-type Ca^2+^ to the SK Ca^2+^-dependent potassium current that shapes the activity profile. Throughout this paper, we consider a fixed level of L-type Ca^2+^ and vary the SK conductance, however it would be equally valid to keep the SK conductance fixed and vary the L-type Ca^2+^. In addition to these two currents, the level of GABAergic inhibition plays a strong role, and we go on to study how it can suppress or sculpt the firing dynamics of DA neurons and their displayed activity modes.

**Figure 1.**
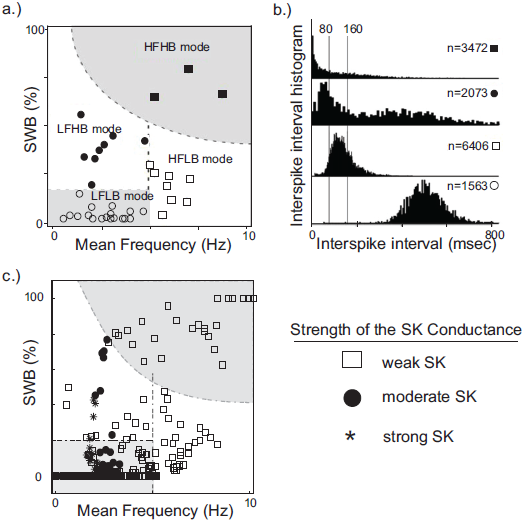
A plot of experimentally observed DA cell firing modes from in vivo rodent VTA adapted from [23] contrasted with data from computational simulations. In (a), the distribution of in vivo firing modes. Along the horizontal and vertical axes are mean firing frequencies and percentage of spikes within bursts, above 20% was considered a high burst neuron. Neurons firing below 5Hz were considered to have a low firing rate, and high firing rate was taken to be neurons firing above 5Hz. In (b), the associated ISI histogram for the data in (a). In (c), we plot the activity modes obtained from considering a family of neurons with varying SK conductances and receiving different levels of drive, *I*_0_ (here GABA is taken to be 0.15). We arrange the data in a similar fashion to [23], but with the shape of the data points indicating the relative strength of the SK conductance (neurons with weak, moderate and strong SK conductance are denoted via the use of squares, disks, and asterisks, respectively). The entire high firing regime is composed of neurons with low SK conductances and a significant portion of the low firing high burst regime are of low and medium SK conductance neurons.

Dopamine neurons receive glutamatergic input from the prefrontal cortex and a combination of glutamatergic and cholinergic drive from the pontine nuclei. Both VTA and SNc are subject to substantial GABAergic inhibition. In particular, the VTA contains populations of GABAergic neurons that provide local GABAergic inhibition to the VTA DAergic neurons (in addition to other GABAergic inputs from the striatum). The VTA GABAergic populations are subject to similar glutamatergic and cholinergic currents as the DAergic populations, and there is an interplay between the DAergic and GABAergic activities. This balance of excitation to inhibition plays a role in the VTA DA neural response to nicotine stimulation [23–25].

Here in our computational study, we find that the degree of GABAergic tone, together with the glutamatergic activation through the NMDA receptors and the intrinsic cellular properties, plays a strong role in shaping a neuron’s firing characteristics. This is not to say that GABA is the most important signal, it appears as though NMDA is likely the principal pathway to initiate burst firing. However, the make-up of the ionic conductances and the degree of the GABAergic tone influence how readily primed a neuron is to burst. So although NMDA can reliably initiate burst firing in DA neurons, the duration of initial burst firing event and the resultant continued firing pattern is dependent on the strength of the DA neuron’s SK conductance.

## Conductance-based DA Neuronal Model

We present a single compartmental conductance-based model for the dynamics of a DA neuron (a schematic shown in Fig. 2) that combines modified conductance mechanisms previously introduced in [11, 15, 26, 27]. Such a model serves to consolidate these approaches to a simplified framework and allows the use of computer simulation to uncover the mechanisms behind the generation of DA activity modes. Parameter values are listed within the text with any unspecified parameters appearing in the appendix.

**Figure 2.**
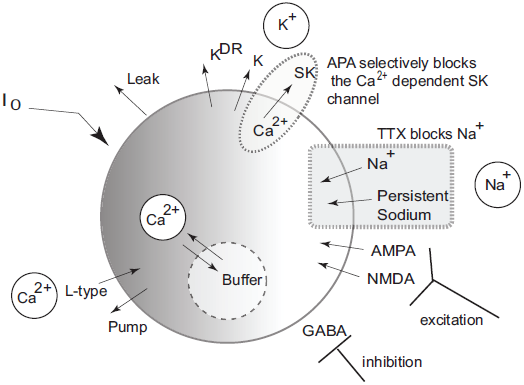
Schematic diagram of the single compartmental conductance-based model of DA neuronal activity. The model includes a generalized intrinsic current whose composition could include in vivo cholinergic drive or in vitro applied currents. The model includes ionic currents for sodium, calcium, and potassium. Both spike generating and persistent sodium currents are expressed, which are both selectively blocked by application of TTX. The potassium currents are composed of a general current, a delayed-rectifier current, an apamin-blocked, calcium-dependent SK-type current. Calcium is predominantly trafficked into the intracellular region via the L-type Ca^2+^ channel and pumped out of the cell via a pump, with the intracellular calcium undergoing fast buffering. The intracellular calcium modulates the SK potassium current and the interaction between the dominant L-type calcium current and the SK potassium current forms the mechanism for an underlying oscillation. In addition to these currents, DA cells receive excitatory synaptic drive through AMPA and NMDA receptors. In addition, DA cells receive inhibitory input from GABAergic neuronal populations.

### i. the membrane potential

The dynamics of the membrane potential are described by

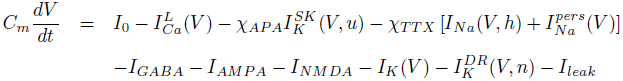

where *I*_0_ is a generic term meant to incorporate the effects due to either an applied current in vitro or a general tonal dendritic input *in vivo*. The L-type calcium current, 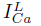, is counterbalanced with the apamin sensitive, calcium dependent SK-type potassium current, 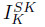 The SK current is dependent on both the voltage and calcium level, with calcium concentration denoted by *u*, and the SK current can be parametrically controlled via the parameter *χ_APA_*. Two tetrodotoxin (TTX) dependent sodium currents are included that can be parametrically regulated via the parameter *χ_TTX_* : (i) a spike-generating sodium current, *I_Na_*, modeled in a Morris-Lecar-like fashion with fast activation, *m_∞_*, and slow inactivation, *h*, (ii) and a persistent sodium current, 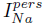 suggested to be of particular import to VTA DAergic neurons [28]. The DA neurons receive GABAergic inhibition and receive excitatory glutamatergic drive through their AMPA and NMDA receptors. We include both a general and a delayed rectifier potassium currents and a general leak current.

A number of observed currents were not included for simplicity. For instance, previous studies have shown that the hyperpolarization-activated cation current, *I_h_*, has little or no effect on VTA pacemaking [28, 29], and hence we omitted it from this study. As a consequence, our model does not reproduce the characteristic depolarization sag after a hyperpolarization event.

### ii. spike generating and voltage dependent currents

The fast processes associated with spike generation involve sodium. The sodium dynamics are taken to be Morris-Lecar-like with fast activation and a slow inactivation gating variable *h*; so that

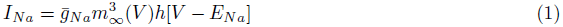

with the conductance 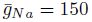 and the reversal potential of *E_Na_* = 55mV. The fast activation is taken to be of the form

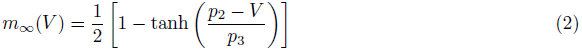

with *p*_2_ = −14mV, *p*_3_ = 11.9, and the inactivation gating variable evolves according to

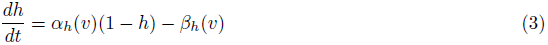

where 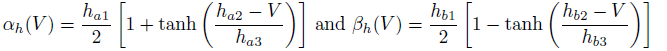

In addition, we include a delayed-rectifier potassium current of the form

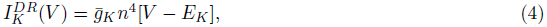

where the conductance is given by 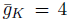 and the reversal potential *E_K_* = −90mV. The associated gating variable evolves according to

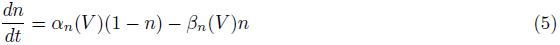

with 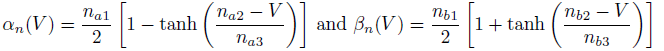

A generic potassium current, *I_K_*, meant to represent an amalgam of the full family of potassium currents, is taken to have the form

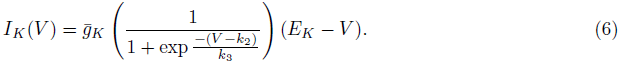

with *E_K_* = −90mV (other parameters listed in appendix).

The model could be further modified to include various Na^+^ currents, such as a TTX independent sodium leak [28]. Here, we included a TTX dependent persistent sodium current [28]. Though not strictly necessary, its presence simplifies parameter selection for the activation variables. The persistent sodium current is taken to be of the form:

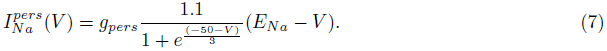

where *g_pers_* = 0.002.

Additionally, we include a general leak current of the form: *I_leak_* = *g_L_*(*E_leak_ − V*) with *E_leak_* = −50mV.

### iii. calcium and calcium-dependent dynamics

The L-type calcium current is taken to have the form

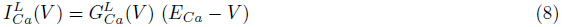

with *E_Ca_* = 100mV and 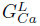 given by,

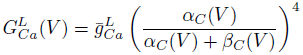

with

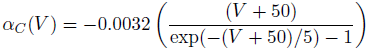

and

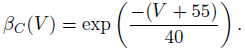

The calcium dependent SK potassium current is given by

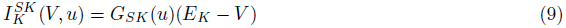

Where 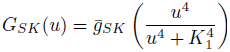

Calcium enters the cell predominantly via the L-type calcium channel (with a minor contribution due to the NMDA channel). The Ca^2+^ is then taken to be ejected via a pump (akin to [27]). The calcium, *u*, then varies according to

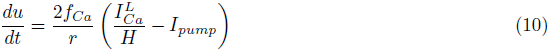

where the cytosolic buffering constant is given by *f_Ca_* = 0.01; *H* is a lumped term involving the valence of calcium and Faraday’s number (*H ≈* 0.0193); the radius, *r*, of the neuron is taken to be *r* = 20*µ*m, and the pump dynamics are taken to be:

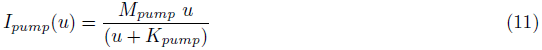

with *M_pump_* = 500 (nM *µ*m)/ms and *K_pump_* = 500nM. Note that since intracellular Ca^2+^ buffering is strong, *u* can be considered a slow variable.

### iv. synaptic currents: GABA, AMPA & NMDA

The two principal regions of the midbrain associated with DAergic neurons, the ventral tegmental area and the substantia nigra pars compacta, both receive substantial amounts of GABAergic inhibition (from the ventral tegmental tail and the substantia nigra pars reticulata, respectively). The GABA cell populations display high frequency, tonic firing patterns and serve to hyperpolarize the target DA cell membrane potential. Since these rates are high, we consider an average conductance parameter for the inhibitory drive, that is, the fine-scale, detailed firing pattern of the GABAergic input is not explicitly tracked, and the current is taken to be:

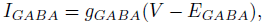

where *g_GABA_* is the conductance and *E_GABA_* = −65mV is the reversal potential. In effect, this current acts akin to a very strong leak current.

The DA cell receives excitatory, glutamatergic drive mediated by AMPA and NMDA receptors. The received AMPA current is taken to be *I_AMPA_* = *g_AMPA_*(*E_AMPA_ − V*), where the reversal potential is *E_AMPA_* = 0 and the conductance evolves according to

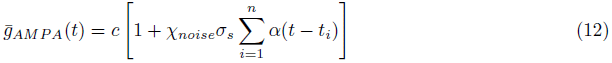

where *c* = 0.002, *σ_s_* is spike strength, and *χ_noise_* is a binary parameter that determines whether noise is included. The spike times *t_i_* are generated via a Poisson process with a rate *λ* (the frequency for the noise typically taken between 10-50Hz). Computational studies with the inclusion of noise produce firing patterns that are more comparable to *in vivo* data that is inherently noisy.

The synaptic influence due to an incoming spike at time *t_i_* is subject to the *α* function:

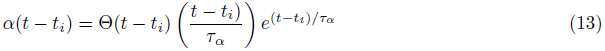

with the time constant given by *τ_α_* = 4ms and Θ represents a heaviside function.

The NMDA current, also, influences DA neuronal dynamics. The NMDA current takes the form:

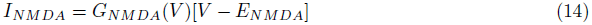

with

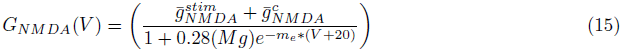

where *M g* denotes the amount of magnesium, taken to be 0.5*µ*M. Here *E_NMDA_* = 0, 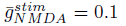 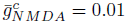 and *m_e_* = 0.08. For simplicity, the parameter 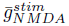 is set to zero, except during NMDA activation / application. The form of the NMDA conductance depends upon the level of magnesium, shown below in Fig. 3, where an increase in Mg^2+^ shifts the conductance curve to be active at higher voltages.

**Figure 3.**
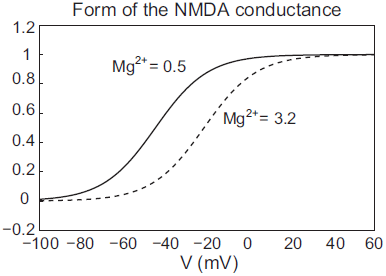
The NMDA conductance depends on both the membrane potential and the level of Mg^2+^. We plot the form of the NMDA conductance for the case of low Mg^2+^(0.5*µ*M) and high Mg^2+^ levels (3.2*µ*M), pictured as a solid and dashed line, respectively.

## Model Results

In the previous section, we presented a pared down representation of a DA neuron that includes an array of observed currents. In the following section, we demonstrate the model’s ability to capture characteristic dynamics of DA neurons. By doing so, we identify conditions for eliciting a variety of firing modes and make a number of experimental predictions. In addition, we examine conditions akin to NMDA bath application [18] for inducing burst firing and conditions akin to DA cell disinhibition (i.e., a release of inhibition from the GABAergic neurons).

### Pacemaking and Burst Firing

Experimentally Ping and Sheperd [22] observed that application of the drug apamin induces burst firing in slice preparations by blocking the SK-channels, which reduces the flow of K^+^, that is, DA cell dynamics depend critically on the strength of the SK-conductance. Motivated by these results, we simulated this experiment by varying the strength of the parameter *χ_APA_*. For *χ_APA_* = 1, this corresponds to a strong SK-conductance and the DA cell fires tonically (Fig. 4a). However as *χ_APA_* decreases, the neuron enters a weak SK-conductance regime and the firing pattern transitions from tonic firing to endogenous or ‘natural’ bursting (Fig. 4b). An initial tight spike doublet occurs at burst onset that is characteristic of DA cell activity [7]. As expected, the strength of the SK conductance strongly influences a DA neuron’s burst firing characteristics.

**Figure 4.**
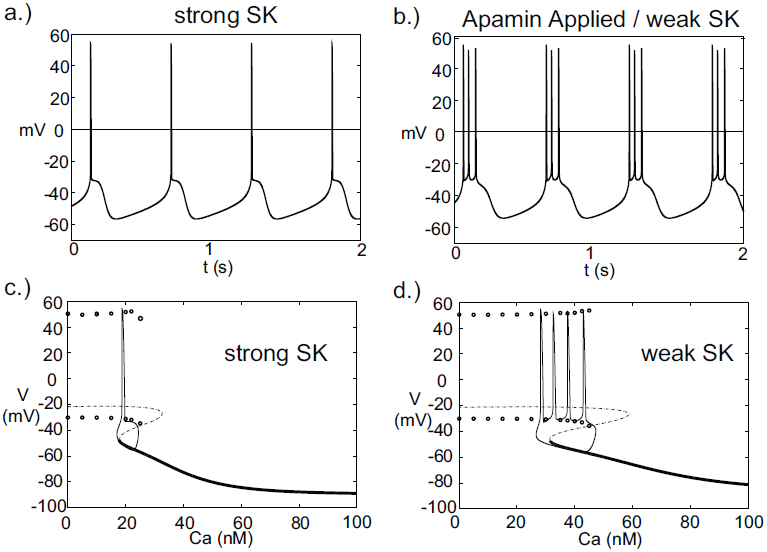
A DA neuron exhibiting a range of firing patterns with a background input of *I*_0_ = 0.2. In (a), the neuron exhibits a tonic firing pattern (*χ_APA_* = 1). In slice preparations, application of the drug apamin suppresses the SK-conductance and yields an increase in burst firing. The mathematical model responds in a comparable manner, by reducing the SK-conductance (*χ_APA_* = 0.2), the DA neural activity pattern enters a burst firing regime (a representative voltage trace is shown in (b)). In (c) and (d), we consider the calcium dependency behind these firing patters by constructing bifurcation diagrams for strong or weak SK conductances, shown in (c) and (d), respectively. We treat the intracellular calcium level as the bifurcation parameter, and as intracellular calcium, *u*, varies from high to low, the hyperpolarized stable steady-state (thick solid line) is lost via a fold bifurcation (with the unstable steady-state branch shown with a dashed line). For low calcium levels, the dynamics enter a periodic orbit and the neurons exhibit fast firing. The extrema values of the oscillations are represented by circles. Trajectories of the full dynamics are superimposed. Note that *I*_0_ = 0 and that the dynamics for a weak SK conductance (*χ_APA_* = 0.2) admit an additional spike during a burst compared to when *I*_0_ = 0.2, as shown in (b).

As the strength of the SK conductance affects DA cell firing patterns and the SK conductance is dependent on intracellular calcium levels, DA neural firing patterns are in turn dependent on intracellular calcium levels. The intracellular Ca^2+^ buffering is strong, rendering *u* a slow variable. We treat intracellular calcium, *u*, as a bifurcation parameter in order understand the firing dynamics over a range of Ca^2+^ levels, see Fig. 4(c,d). Consider 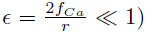 as 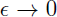 the intracellular calcium level becomes approximately fixed. For ‘large’ levels of calcium, the membrane potential attains a stable hyperpolarized steady state. As *u* decreases, the stable steady-state is lost through a fold bifurcation. For ‘low’ Ca^2+^ levels, the neuron enters a fast firing regime. This makes intuitive sense as for low Ca^2+^, the SK-channel remains inactive, while for high Ca^2+^, the SK-channels remain open resulting in a hyperpolarized state due to potassium leaving the cell in a non-gated fashion. When the full dynamics are included, starting from a hyperpolarized state, the neuron slowly becomes depolarized; simultaneously, the intracellular calcium level slowly decreases. Eventually the trajectory, passes the fold bifurcation and enters a periodic orbit associated with neural firing. In this depolarized periodic orbit, intracellular calcium levels rises. When intracellular calcium levels rises sufficiently to activate the SK potassium current, the trajectory falls off of the periodic orbit and returns to a hyperpolarized ‘resting’ state. The activation of the SK-current coincides with the periodic branch colliding with the unstable steady state branch at a SNIC bifurcation. The shape of the bifurcation diagram is dependent on the relative strength of the SK-conductance (*χ_APA_*). For strong SK-conductances, the neuron fires once (twice in some instances) before returning to a rest state, whereas for weak SK-conductance, the ‘bend’ in the bifurcation diagram is more pronounced and there is a wider range of calcium levels where the periodic orbit occurs before colliding with the steady state branch. As a result, for weak SK conductance, the neuron goes through a greater number of oscillations in that depolarized oscillatory orbit before falling off the orbit and returning to the hyperpolarized state, i.e, the interburst quiescence. In addition, the interspike intervals (ISIs) increase through burst duration as is indicative of classical square-wave-like dynamics [30]. In future studies, we will be further examining the structure of the dynamics by performing fast/slow analyses with averaging.

### Depolarization Block

The introduced model for DA cell activity undergoes depolarization block when overly driven, and we construct bifurcation diagrams to study the role of the amalgam term, *I*_0_, that represents contributions due to an applied current or bulk drive to the neuron, see Fig. 5. For small values of *I*_0_, the neuron (for both weak and strong SK conductances) has a stable hyperpolarized steady-state, and the neuron exhibits quiescence. As *I*_0_ increases, the steady-state loses stability through either a subcritical Hopf or through a fold bifurcation for strong and weak SK-conductances, *χ_APA_* = 1, 0.2, respectively. As *I*_0_ increases, the frequency of periodic orbits increases, that is, the neuron fires with an increasing frequency. For sufficiently large *I*_0_, the periodic orbit is lost and the neuron enters as stable steady-state, associated with depolarization block.

**Figure 5.**
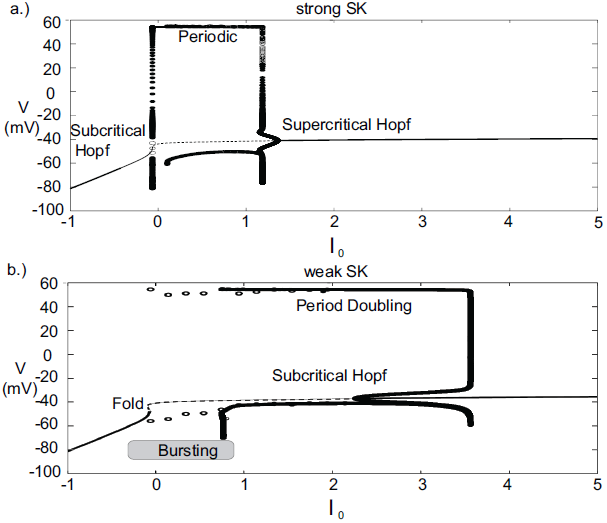
Bifurcation diagrams for understanding how DA firing patterns depend upon the variable *I*_0_ for strong or weak SK conductances, (that is, *χ_APA_* = 1.0 and *χ_APA_* = 0.2), shown in (a) and (b), respectively. Here we plot membrane potential with the generalized intrinsic / applied current variable *I*_0_ taken as the bifurcation parameter. In both cases (weak and strong SK conductance), for small *I*_0_, the neuron exhibits quiescence. For increased *I*_0_, the dynamics follow a periodic orbit, i.e., the neuron fires repetitively. For larger *I*_0_, the neuron undergoes depolarization block. The level of *I*_0_ required to reach depolarization block depends inversely on the strength of the SK conductance. Note that for weak SK conductance for *I*_0_ between approximately 0 and 1.0 that the periodic orbit displays a mixed-mode oscillation, indicative of endogenous burst firing, as *I*_0_ increases, the firing displays ‘fast’ pacemaker firing. Stable steady states are shown by thick lines and periodic solutions are labelled.

For weak SK conductance (*χ_APA_* = 0.2) and the same progression from low to high levels of *I*_0_, the neuron begins at a hyperpolarized steady-state and enters a burst firing mode, that is a mixed mode oscillation at *I*_0_ *≈* 0. As *I*_0_ further increases, the frequency of the burst firing increases. At *I*_0_ *≈* 1, the burst firing pattern transitions to a fast firing tonic pattern, where the mixed-mode oscillations appear to be lost. Past *I*_0_ *≈* 3.5, neurons exhibit depolarization block and attain a stable depolarized steady-state solution. The exact nature of the substructure of the periodic firing patterns displayed for weak SK neurons and the transitions between the mixed-mode to fast firing is not examined in this work but is to be the subject of a future study. Beyond exhibiting multiple activity modes, a comparison of the bifurcation diagrams for strong and weak SK conductances shows that neurons with SK conductance have periodic orbits (i.e. fire) for a wider range of *I*_0_. This suggests that neurons with weak SK-conductances are more resistant to depolarization block, that is, neurons with a weak SK conductance will require higher levels of *I*_0_ before a cessation of firing. Moreover, neurons with an extremely weak SK conductance and receiving high levels of drive may enter a tonic fast firing regime. A strong SK conductance appears to be the key to property of depolarization block observed for many DA neurons.

### Burst termination via depolarization block

The model exhibits a wide range of firing modes / behavior dependent on the make-up of an individual neuron’s ionic currents, to say nothing of the effects of differences in broader system level activity changes in the neural circuit. For DA cells, classic burst firing is initiated by a spike doublet followed by intraburst spikes with ISIs that are approximately equal as classified in [8]. However, burst firing patterns of a markedly different structure have been observed, e.g., even in the same Grace-Bunney paper [8]. Some of the burst firing patterns display decreasing spike amplitudes and increase of firing frequency during the burst event, so that the ISI decreased within a burst. These burst events appear to terminated via depolarization block. This type of bursting pattern was also found in the classic paper of Ping-Shepherd [22] in SNc DA neurons in slice preparations under apamin application and has been viewed in VTA extracellular recordings [20, 21].

It is regarded as a characteristic of DA neurons that they are susceptible to depolarization block, whereby an increase of an applied current results in a cessation of DA cell firing due to sodium inactivation. In this work, we have identified an alternate set of parameters where individual neurons exhibit depolarization block in their endogenous dynamics, that is, the bursts terminate via depolarization block. The changes to the parameter regime include a broadening of the region of sodium activation, strengthening and altering the delayed-rectifier potassium current, and changing the dynamics of the sodium inactivation gating variable. The detailed changes of parameters are listed in the appendix.

With these parameter changes to our model, the observed dynamics display the following described behaviors. For moderate levels of *I*_0_, neurons with a strong SK conductance exhibit regular firing at relatively high frequencies for DA neurons. However, for the same level of *I*_0_, weak SK-conductance neurons exhibit endogenous burst firing where the bursts are terminated via depolarization block, shown in Fig. 6(a,b). Moreover, both the spike amplitude and ISI decreases throughout the burst duration, and burst termination coincides with a prolonged depolarization plateau. We plot these trajectories in phase space with respect to the membrane potential, *V,* and sodium inactivation, *h*, see Fig. 6(c,d). We note that for a weak SK conductance (*χ_APA_* = 0.2), a neuron, initially at a hyper polarized state (labeled H), becomes slowly depolarized and then enters a burst firing event with 4 spikes. During this burst event, both the spike amplitudes and the ISI decrease, that is, the spikes get smaller and there is an increase in the frequency during the burst. At the end of the burst event, the trajectory is at the point labelled D. Here the sodium inactivation is small and the trajectory lingers at the intersection of the *V* and *h* nullclines, an unstable steady state associated with long the plateaus of depolarization.

**Figure 6.**
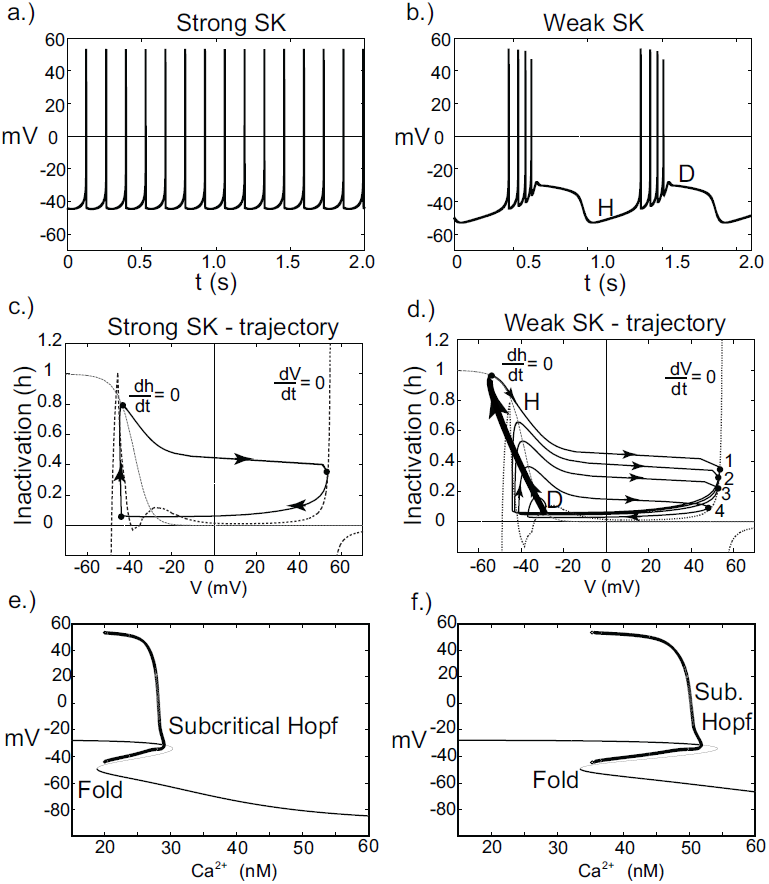
Instances of DA neuronal bursting terminated via depolarization block. In (a), we observe that for these parameters and for a strong SK-conductance (*χ_APA_* = 1), this model DA neuron exhibits a relatively fast (approximately 7 Hz) tonic firing pattern. Contrast this regular firing pattern with the burst firing that occurs for a weak SK-conductance (*χ_APA_* = 0.2), shown in (b). For weak SK conductances, the DA cell exhibit endogenous burst firing where the bursts are terminated by depolarization block via the intrinsic rhythm of the neuron. In (c,d), we plot the associated trajectories the membrane potential V and sodium inactivation h. Nullclines for both V and h are included (see labels). In (d), we note the precipitous drop in h during a burst and the termination of the burst (at point D) is an unstable depolarized state, associated with the plateaus of depolarization shown in (b). In (e) and (f), we show corresponding bifurcation diagrams for (a) and (b) taking the intracellular Ca^2+^ level is treated as the bifurcation parameters. As intracellular calcium, u, varies from high to low, the hyperpolarized stable steady-state (thick solid line) is lost via a fold bifurcation (with the unstable steady-state branch shown with a thin line). The trajectory then enters a periodic orbit whereby the neurons are said to be firing. The extrema values of the oscillations are represented by circles. For low calcium levels, there is a stable depolarized steady-state solution.

In this modified parameter regime, Ca^2+^ buffering is fast, rendering Ca^2+^ dynamics slow. We take the intracellular calcium to be at quasi-steady state and generate a bifurcation diagram in order to better understand the dynamics of the full system, shown in Fig. 6(e,f). The bifurcation structure contains a fold and a subcritical Hopf. For the case of a strong SK conductance, a hysteresis loop for regular firing exits whereby a single spike occurs followed by entering the hyperpolarized state. As Ca^2+^ decreases, the trajectory falls from the fold and another spike occurs. For low levels of Ca^2+^, there exists a stable depolarized steady-state. In the case of a weak SK conductance, the shape of the bend in the curve is more pronounced and there is a larger regime for the periodic branch (this is an oscillation in voltage and corresponds to neural firing). As the neuron fires, Ca^2+^ ramps up in the cell leading to the activation of the SK current. As this occurs on the bifurcation diagram, the trajectory would be approaching the subcritical Hopf bifurcation from left to right. When passing this bifurcation, the trajectory lingers on the unstable (depolarized) fixed point before dropping to the lower stable branch (i.e., the hyperpolarized state).

As the neuron loses Ca^2+^, the neuron becomes slowly depolarized. Eventually, it loses the stable steady-state via a fold bifurcation and enters the periodic orbit again, associated with neural spiking. During this spiking stage, intracellular Ca^2+^ rises and the trajectory moves to the right in the bifurcation diagram. As this occurs, the amplitude of the spikes decreases during the burst progression, as also do the ISIs (that is, the spike frequency increases during a burst event). At burst termination, the trajectory lingers at the unstable depolarized steady state before ‘falling’ down to the hyperpolarized interburst steady-state.

Another difference for this parameter choice is that both the strong and weak SK conductance neurons have similar depolarization block properties with respect to the parameter *I*_0_, as seen in Fig. 7. Both strong and weak SK conductance neurons are more impervious to increases in *I*_0_.

**Figure 7.**
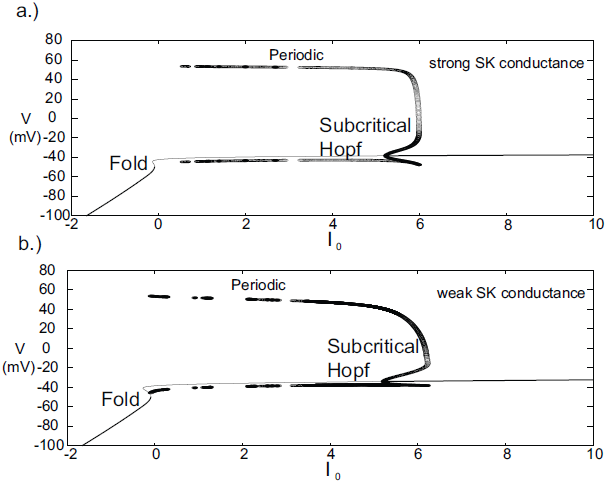
Bifurcation diagrams for an alternate parameter set treating *I*_0_ as a bifurcation parameter. Here we consider both strong and weak SK conductances with *χ_APA_* = 1.0 and *χ_APA_* = 0.2 shown in (a) and (b), respectively. Note the presence of both a Poincaŕe-Hopf-Andronov bifurcation and a saddle-node on an invariant circle (SNIC) bifurcation. Also, a far greater *I*_0_ is needed in order to induce depolarization block, and this value of *I*_0_ is similar for both strong and weak SK conductances. Stable steady states are shown by thick lines and periodic solutions are labelled.

### Quantifying Bursts

The seminal work of Grace and Bunney [7] examined the properties of in vivo bursting and introduced criteria for identifying the bursts. They noted that burst onset was associated with a tight doublet of two successive spikes, and thus identified burst onset by “the concurrence of two spikes with an interspike interval (ISI) of less than 80ms. Burst termination was defined as an interspike interval of over 160ms.” This is a straightforward, algorithmic definition of a burst firing event and is still widely utilized. With this definition, one often employs a measure of the degree of burst firing that corresponds to the percentage of spikes occurring within bursts (spikes within bursts, SWB). Due to its prevalence in the literature, we consider this familiar burst measure (SWB), however with this definition of bursting with increased neuronal activity, it classifies fast, tonic firing to be equivalent to a burst of an extremely long duration. Thus, we contrast the SWB measure with an alternate burst measure introduced by Van Elburg and Van Ooyen [31].

In order to differentiate, high rate, tonic firing from truly ‘bursty’ dynamics, we consider the Van Elburg / Van Ooyen measure for degree of bursting. This measure utilizes both the interspike and ‘two-spike’ interval. Take ISI to be the distribution of the interspike intervals over an experiment. In addition, let TSI represent the ‘two-spike interval’ distribution for the same spike train, which is made up of time intervals corresponding to the time between a spike and the second subsequent spike. If *σ_I_* and *σ_T_* represent the standard deviation of the ISI and TSI distributions, respectively, and *µ_I_* is the average value of the ISI distribution. The Van Elburg / Van Ooyen measure for the degree of bursting is given by

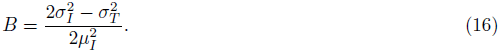

Using this measure, *B* above a threshold value, *B*_Θ_, would constitute a bursting neuron. For example, in [31], a neuron with a measure *B > B*_Θ_ = 0.15 was considered to be bursting.

### DA Activity Modes

A wide range of parameter values were utilized in order to examine the necessary conditions to elicit certain modes of activity that have been observed by [23, 32] and others. Other laboratories are working to identify (and classify) DA neurons from electrophysiological properties [33–36]. Here we consider a range of strengths of the SK conductance and a range of values for the level of the intrinsic / background drive variable *I*_0_, shown in Fig. 8, in order to study the mechanisms underlying multiple activity modes. From the phase diagram, we deduce that for moderate to strong SK conductance strengths, the DA neurons undergo depolarization block when given a large applied current, *I*_0_. The model results suggest the experimental prediction that neurons that have undergone apamin exposure or that have an inherently lower SK conductance will be more resistant to depolarization block (that is, require a greater applied current to initiate it). There appears to be an inverse relationship between SK-conductance strength and the needed applied current to bring on depolarization block. In addition, neurons that display higher firing rates appear to be correlates with lower SK levels.

**Figure 8.**
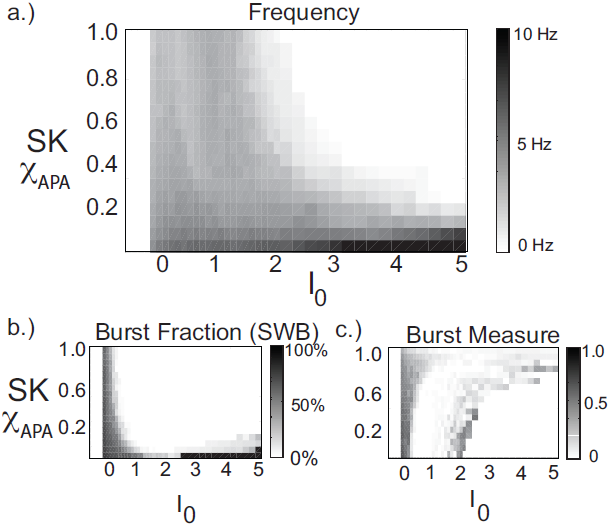
In these phase diagrams, we show how firing frequency, burst fraction (SWB), and the burst measure depend on both *I*_0_ and the strength of the SK-conductance influence DA cell firing patterns. In (a), the firing frequency is plotted with *I*_0_ and the strength of SK-conductance varying and receiving noise in the AMPA conductance at a rate of 50 Hz. In (b) and (c), the level of bursting is similarly plotted using the classic percentage of spikes within bursts (b) and the burst measure proposed by [31]. For small *I*_0_ the system displays quiescence. For “moderate” *I*_0_, DA cells display activity (neurons with a lower SK conductance fire at a moderately higher rate). For larger *I*_0_, the DA cells cease firing (that is they undergo depolarization block) and the level of *I*_0_ required to initiate depolarization block is inversely proportional to the SK-conductance. That is, neurons with a low SK-conductance continue to fire at levels of *I*_0_ that would cause a neuron with a high SK conductance to cease firing.

By considering the two measures of bursting (SWB measure and the Van Elburg/Van Ooyen measure), underlying differences in the firing patterns can become apparent. The SWB measure classifies neurons with a weak SK conductance that fire tonically at approximately 8 Hz to be bursting, whereas the burst measure of Van Elburg and Van Ooyen does not classify these neurons as bursting. Moreover, the Van Elburg and Van Ooyen measure accentuates the presence of mixed-mode oscillations within the firing patterns that are associated with burst firing. Another difference between the measures is that at a level of *I*_0_ where neurons are undergoing depolarization block, the neurons fire sporadically due to noise. At these points, the burst measure classifies these neurons to be bursting.

The strength of the SK conductance and the intrinsic/applied current do not alone determine a DA neuron’s activity mode. The level of GABAergic drive plays an import role in determining the activity profile of a DA neuron as noted in [37–39]. We generate a series of diagrams akin to [23] to study the impact of GABA on influencing DA activity modes (we consider low, medium and high levels of GABA), shown in Fig. 9. The most telling conclusion that we infer from these plots is that the strength of the GABAergic tone plays a dominant role in DA cell activity mode determination. For a strong GABAergic tone, burst firing is reduced and the activity modes are more tonic in nature. As GABAergic tone decreases, firing rates increase as well as the number of neurons exhibiting burst firing. This suggests that if the nAChRs within the GABAergic population are exposed to nicotine and enter a desensitized state, the resulting disinhibition of the DA cell population will result in a bulk increase of the average DA cell activity, but also an increase in burst firing. This increase in bursting occurs predominately in neurons that have weak SK conductances, so this supports the hypothesis of [23] that neurons that display some degree of bursting are neurons that display increased burst responses to nicotine simulation, that is, there are neurons that are primed to burst. One matter to note is that with either measure of bursting, spike doublets are classified as bursts and much of the ‘low firing, high burst’ activity are, in fact, doublets generated periodically.

**Figure 9.**
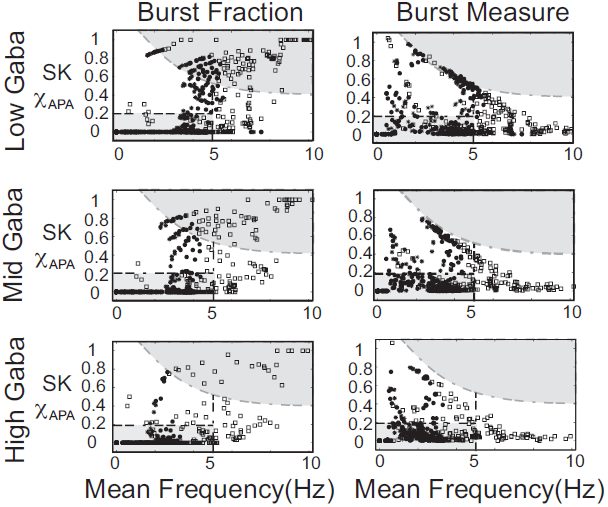
The strength of the GABA inhibition has a strong influence on the firing and burst firing characteristics of a DA neuron over a range of neuronal drive levels and SK-conductance strengths. Neurons with weak, moderate and strong SK conductance are denoted via the use of squares, disks, and asterisks, respectively. Here GABA levels listed as low, mid, and high corresponds to *g_GABA_* = 0.01, 0.02, and 0.03, respectively. The GABA conductance included is taken to be constant and does not capture the dynamic network level changes in the mean field activity of the GABA neuronal population innervating the DA neuron.

Differences in the degree of GABAergic control to a DA neuron greatly influences the resulting firing pattern. In rats under anesthesia via chloral hydrate, only 18% of the observed DA neurons in the SNc exhibited bursting, whereas, 73% of the VTA DA neurons exhibit burst firing [40]. In our numerical studies, we find that neurons receiving more GABAergic inhibition exhibit more pacemaker-like firing, which is more indicative of SNc DA neurons. Thus, we postulate that the degree of GABA inhibition to the SNc DAergic neurons is greater than that to the VTA DAergic neurons.

### A Possible Novel Firing Mode Unveiled

In our numerical studies, numerous phase diagrams were generated considering many sets of parameters and amounts of noise in the glutamatergic synaptic drive. For a wide range of parameters, phase diagrams akin to the one pictured in Fig. 9 were obtained. However, we also discovered another parameter set (listed in the appendix) that exhibited a number interesting characteristics. The associated phase plots for this parameter set are shown in Fig. 10, where the activity modes for a number of GABA levels are considered as well as a range of SK conductances and values of *I*_0_.

**Figure 10.**
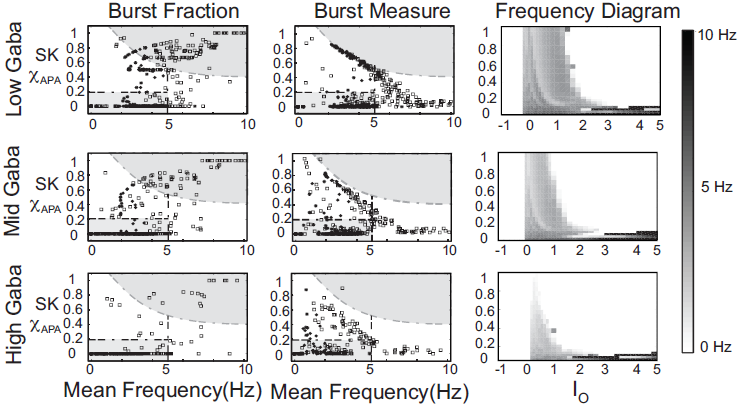
An alternate phase diagram that suggests a novel activity mode of DAergic neurons. The strength of the GABA inhibition has a strong influence on the burst firing characteristics of a DA neuron over a range of neuronal drive levels and SK-conductance strengths. Neurons with weak, moderate and strong SK conductance are denoted via the use of squares, disks, and asterisks, respectively. Again, GABA levels listed as low, mid, and high corresponds to *g_GABA_* = 0.01, 0.02, and 0.03, respectively.

We observe that there is a pronounced sensitivity to the level of GABAergic inhibition, whereby for high GABA levels, much of the neuronal activity is silenced with the burst firing modes almost completely removed. Furthermore, the threshold for *I*_0_ to induce depolarization block varies greater at the different levels of GABA inhibition. Akin to the previous phase diagram, the firing rate of the neurons increases with a decrease of the SK conductance. However, the “very high” firing rates plotted here not only rise above 10Hz, but as *I*_0_ raises past 3, the firing rates enter fast firing regimes between 20-40Hz. Such a tonic, fast firing regime is interesting for a number of reasons. For moderate levels of GABAergic inhibition (Mid GABA), for neurons with low SK conductance (0.15 *< χ_APA_ <* 0.2), as *I*_0_ increases, these neurons cease firing but with a further increase of *I*_0_, these neurons enter a fast firing regime. This fast firing regime may occur for extreme parameter values, but it may signify a subclass of DAeric neurons that defy the convention that DAergic firing rates are less than 12Hz, as has been suggested by the findings of Lammel et al. [35]. These computational findings taken together with the data of [35] suggest the possible existence of a novel activity mode yet to be unveiled and yields to the experimental prediction that neurons exposed to apamin may display similar physiological dynamics. In Fig. 11, we go onto examine the mathematical structure of this novel firing mode. We observe the emergence of a periodic solution, i.e., very fast neural firing.

**Figure 11.**
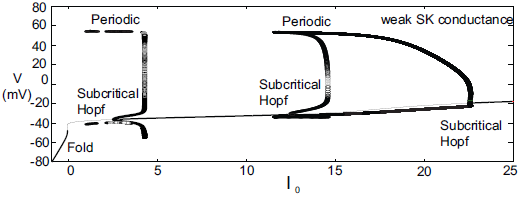
Bifurcation diagram for moderate GABA level with a weak SK conductance (*χ_APA_* = 0.2) to study the neural dynamics over a range of *I*_0_. The observed results are not in perfect accord to the phase diagram shown in Fig. 10 for the frequency distribution at moderate GABA level as there is no inclusion of noise in this diagram. However we observe that as *I*_0_ increases from 0, the neuron enters a firing regime, but as *I*_0_ increases further it enters a state of quiescence. With a further increase of *I*_0_ the neuron can enter another fast firing regime. Stable steady states are shown by thick lines and periodic solutions are labelled.

### NMDA-Induced Firing

Other than endogenous burst firing behavior, increases in the NMDA conductance elicits transient or endogenous burst firing. This is inline with the observed endogenous bursting patterns in vitro [18] due to NMDA-bath application and is consistent with the NMDA-initiated burst firing in vivo [41]. Li et al. [42] first modeled NMDA-induced bursting using a two-compartmental representation of the the neuron, which relied on somatodendric interactions in order to generate burst firing. Here, we utilize the original parameter set in our single-compartment model (with sodium activation slightly altered, *p*_3_ = 12.5, though these properties are general over a range of different parameters). We consider a range of SK conductances and different levels of Mg^2+^(different levels of Mg^2+^ alter the activation dynamics of the NMDA current, see Fig. 3). In Fig. 12a, a DA neuron with a ‘strong’ SK conductance and relatively low Mg^2+^ level, transitions from a tonic firing regime to a transient burst of activity at time 2.0s by an instantaneous increase of the NMDA conductance, *g_NMDA_*, by 0.1ms/cm^2^ for a duration of 2.0s. After the burst coincides with the onset of the stimulus (sudden rise of the NMDA conductance) and then enters a depolarized state. (Note that whether the neuron enters a fully depolarized state depends upon the variables associated with sodium activation and inactivation). At the end of the stimulus, the neuron becomes hyperpolarized and displays a long recovery period before returning to pacemaker-like firing. For the same level of Mg^2+^(0.5*µ*M), a DA neuron with a weaker SK conductance can be similarly be induced to burst via an increase of the NMDA conductance (see Fig. 12c). The resulting burst firing event is of a longer duration (that is, “burst firing is augmented, not suppressed, by SK channel blockade” [14] and here is followed by pacemaker-like firing for the duration of the increased NMDA level.

**Figure 12.**
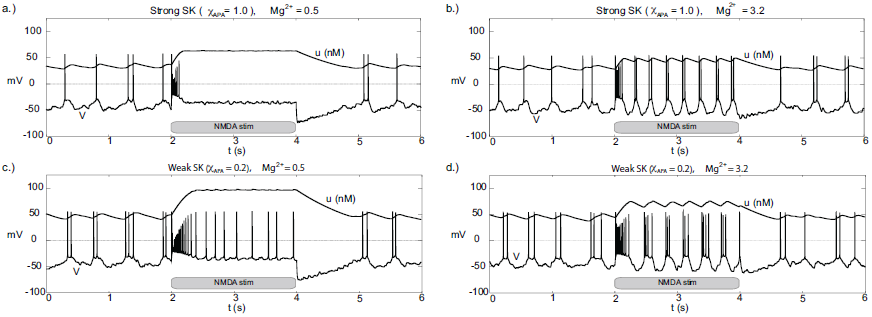
Differential burst firing events due to an increase in the NMDA conductance for *I*_0_ = 0.3, responses depend on both the strength of the SK conductance and the Mg^2+^ level. In (a), a neuron with a relatively strong SK conductance (*χ_APA_* = 1.0) has NMDA applied at time *t* = 2*s* (manifested by an immediate rise of the NMDA conductance), which elicits a a burst firing event. In (b), the preparation is repeated but with a weak SK conductance (*χ_APA_* = 0.2), results in a significantly longer burst firing event followed by pacemaker-like firing for the duration of the stimulus. In (a,b) the Mg^2+^ level was 0.5, in (c,d), we repeat the experiment at an elevated level of Mg^2+^ and note the resultant endogenous burst firing during stimulation (akin to [18]). In addition to tracking the membrane potential, we also display the intracellular calcium dynamics. These simulations include noise in the AMPA drive of 40Hz.

For a similar rise in the NMDA conductance but with high levels of Mg^2+^(3.2*µ*M as in [18]), at stimulus onset an initial burst event is induced (whose duration depends on the strength of the SK conductance). Following that initial burst, the neuron enters a burst firing regime (the degree of bursting dependent on the strength of the SK conductance) as shown in Fig. 12(b,d). In effect, with high levels of Mg^2+^, the NMDA conductance varies more during the observed rhythm. When the neuron is in a ‘down’ state, the NMDA conductance is small and this accentuates the rhythm in part due to the interaction between the L-type Ca^2+^ and the SK-current. Moreover, during the plateaus of depolarization the NMDA conductance is active. We note that in the low Mg^2+^ regime (shown in Fig. 12(a,c)) that the NMDA current is present and driving the neuron, and with the rise in the conductance level, this NMDA current acts akin to a rise in *I*_0_. Please note that even for neurons receiving high levels of GABA with completely suppressed activities that NMDA can still initiate transient burst events (figure not shown).

Recall that the phase diagrams suggest that SK-weak DA neurons require higher levels of *I*_0_ before they undergo depolarization block. These results suggests that the neurophysiological response to an NMDA application can be used to differentiate weak versus strong SK DAergic neurons, that is, the relative strength of the SK conductance could be elucidated physiologically via differential excitable responses. Our finding suggests that in addition to weak SK neurons displaying a wide range of endogenous burst firing regimes, they also respond to stimuli in a more ‘bursty’ fashion. This is line with the observation of [23], that in response to nicotine, initially bursting neurons are more apt to enter stronger burst firing regimes, that is, it is unlikely that initially tonic firing neurons will enter burst firing regimes.

We consider the neural response to bulk rises in the NMDA conductance by DA neurons receiving a strong applied current (that is, we consider a relatively large value for the parameter *I*_0_ = 2.5). For both low and high Mg^2+^, when the drive is high *I*_0_, neurons with a strong SK conductance are near depolarization block (with sporadic firing due to noise) and neurons with a weak SK conductance exhibit ‘fast’ pacemaker firing, see Fig. 13. For low Mg^2+^(Fig. 13(a,c)), an instantaneous rise in the NMDA conductance at *t* = 2*s* elicits a similar response to the experiment performed with moderate *I*_0_ (*I*_0_ = 0.3). Akin to the previous simulation at moderate levels of *I*_0_, at the completion of the stimulus the cell enters a hyperpolarized state. By and large, there is a decrease of activity due to the prolonged raise in the NMDA levels. For higher levels of Mg^2+^, during the period of elevated NMDA, the neurons display fast firing (that is, elevated NMDA elicits an initial burst then leads to an increased firing rate for both weak and strong SK conductances, Fig. 13 (b,d). For high levels of Mg^2+^, the degree of hyperpolarization following the NMDA stimulus is diminished.

**Figure 13.**
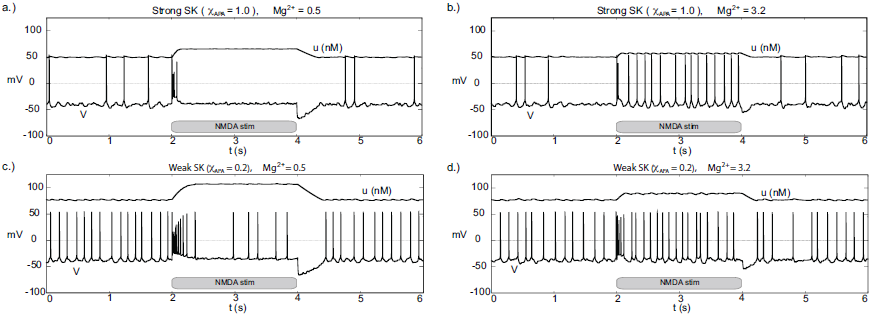
Burst firing events from depolarized (or near depolarized) states due to NMDA increases for neurons of strong and weak SK conductance (*χ_APA_* =1.0 and 0.2) shown in (a) and (c) for low Mg^2+^ levels, respectively. NMDA increases by 0.1 at time *t* = 2s for a duration of 2s. In (a), there is a transient 4 spike burst, while for a weak SK conductance (shown in (c)) we observe a prolonged 9 spike burst event followed by sporadic activity until the NMDA stimulus is released. In both instances, at the release of stimulation, there is a pronounced hyperpolarization of the membrane potential. In (b,d), we consider similar conditions but for high levels of Mg^2+^. We find that in each case (for both weak and strong SK conductance) that the rise in the NMDA conductance at time *t* = 2s results in quasi-regular firing.

### Disinhibition-Induced Firing

Through an examination of the phase plots of the activity modes for various levels of GABA, we noted the profound effect of GABA on influencing the firing modes. However, that study was for the endogenous firing patterns of neural dynamics. Here we consider transient disinhibition events (akin to [39]). From a state that exhibits low neural firing due to the degree of high GABAergic tone), a release from inhibition results in transient burst firing event followed by repetitive firing in a slow tonic or burst firing pattern, dependent on the the SK-conductance of the neuron, Fig. 14. The simulated disinhibition event was represented via changes in the conductance level for the GABA, where high GABA levels transition to low GABA during the disinhibition event, that is, *g_GABA_* = 0.04 *→* 0.01) during the disinhibition event, that is, this disinhibition represents a decrease in the shunting current, rather than an increase in a constant drive current, say *I*_0_. One difference that we note is that after a disinhibition event there is no resulting hyperpolarization as found after an NMDA bath application. In addition, the model results differ from those of [39] where they attain prolonged periods of increased fast firing burst events. Here the burst events differ. An initial burst firing event occurs at disinhibition onset followed by a modulated firing pattern of the original firing pattern (which may display of degree of bursting dependent on the SK conductance of the neuron, see Fig. 14c). For the parameters considered here, for high Mg^2+^ levels with strong inhibition was near quiescent, so that neural activity was relegated to the period of disinhibition (see Fig. 14(b,d)) as the NMDA current would contribute very little with Mg^2+^=3.2*µ*M and *V* ≈ −50mV (please refer to Fig. 3).

**Figure 14.**
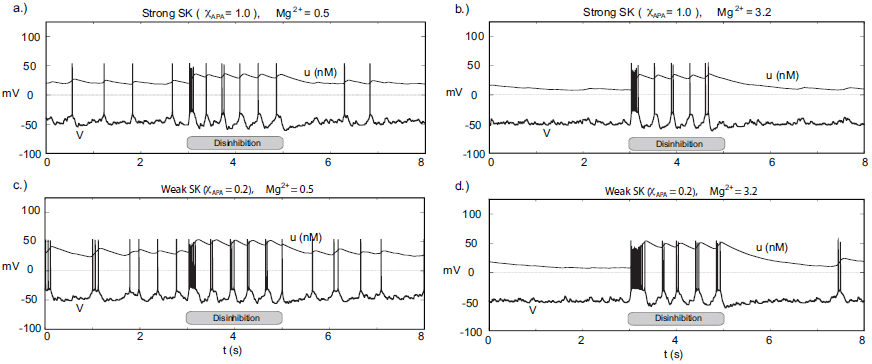
Transient disinhibition can induce burst firing in a manner dependent upon the strength of the SK conductance and the level of Mg^2+^(here *I*_0_ = 0.3). From an initial state of high GABA inhibition, g*_GABA_* = 0.04, where each neuron displays little neural activity, with a release from inhibition (*g_GABA_* = 0.04 → 0.01) at *t* = 3, all the neurons display a transient burst firing event at the onset of disinhibition (whose duration depends on the SK conductance and the level of Mg^2+^). All neurons display an increased firing rate during the period of disinhibition and display either pacemaker-like or burst-like firing dependent upon the strength of the SK conductance. Following the period of disinhibition, the cell enters returns to its previous firing pattern with no appreciable post-stimulus hypepolarization.

## Discussion

Here in this paper, we have utilized computational modeling in order to better understand the mechanisms controlling the ensemble of DA neuronal classes found in DA neurons: both endogenous rhythms and stimulus-induced firing patterns. These findings are in line with experimental findings and complement previous theoretical modeling efforts, however in a single compartment framework as somatodendritic interactions appear to not be a necessary for burst firing initiation [13, 14] (in particular the distal dendrites).

This work accentuates the importance and roles of the SK conductance, GABA inhibition, and the NMDA current have in burst generation. Our results suggest that neurons with a relatively low SK conductance fire in a fast firing mode, are correlated with burst firing, and require higher levels of applied current before undergoing depolarization block. Due to the interplay between the L-type Ca^2+^ and the SK conductance, another way to consider these low SK conductance neurons would be strong L-type Ca^2+^ neurons. However, a decrease of the SK conductance is of potential physiological significance since such slow K-channels are the primary target for down-modulation by ACh through muscarinic AChRs (mAChRs), hence ACh action through mAChRs may prime the DA neurons for bursting (e.g., [43]). Further importance of the potassium currents is evidenced by an alteration of the potassium dynamics, which we found can lead to neural bursting that terminates via depolarization block, akin to firing pattern observed in vivo in VTA DA neurons and endogenous burst dynamics in SNc DA cell preparations under apamin application. The spikes within these bursts are of decreasing amplitude and have decreasing ISIs. This suggests a markedly different bifurcation structure behind the dynamics. Due to the existence of two characteristic burst firing patterns, we postulate that this may be indicative of VTA DA bursts arising due to two different methods: NMDA driven versus a release from GABA inhibition.

Our computational studies on NMDA and GABA are in line with the findings of [37] in the context of multi-compartmental models and with the study of [39]. The firing patterns we observed were characterized by an initial burst event and then enters either a firing mode dependent on the characteristics of the cell (that is, not a sustained high frequency firing for the duration of the disinhibition). The model reinforces the framework that NMDA levels, strength of the SK-conductance, and the level of GABAergic inhibition are all important components in determining the firing mode of a DA neuron, however it further refines the roles of the individual components. The level of GABAergic inhibition strongly influence the endogenous burst firing properties of neurons, that is, strong GABA inhibition leads to mostly tonic firing activity modes. Whilst a removal of that inhibition, i.e., disinhibition, can induce transient burst disinhibition events. Moreover, the main initiator of burst firing events is likely NMDA. We can differentiate NMDA stimulation versus disinhibition events by the subsequent drop in the membrane potential after prolonged NMDA stimulation.

The activation properties of the NMDA receptors can dramatically affect the response properties of neurons. In our studies, we considered a range of NMDA conductances and vary the amount of Mg^2+^. Note that references to high or low amounts of Mg^2+^ are equivalent to shifting the activation profile for the NMDA receptors (see Fig. 3). In NMDA stimulation experiments, DA neural response was characterized by a transient burst event followed by either quiescence or pacemaker firing in the case of low Mg^2+^, whereas for high Mg^2+^, DA neurons displayed pronounced bust firing or high firing rates. The response properties of NMDA stimulation differed from disinhibition events whereby neurons displayed an initial burst at disinhibition onset followed by either pacemaker-like or a burst firing for neurons with strong and weak SK conductances, respectively. Taken together, the above results suggest to obtain strong endogenous bursting under NMDA stimulation, neurons should have both weak SK-conductances and high levels of Mg^2+^, whereas for disinhibition events, the strength of the SK conductance alone appears to account for endogenous burst firing patterns.

In considering the role of the SK conductance and GABAergic tone, we identified a set of parameters whereby neurons with weak SK conductance could undergo a cessation of neural firing via an increase in *I*_0_, however with further increases of *I*_0_ would enter a fast spiking firing mode. This puts into question the ubiquitous belief that DA neurons fire at a rate less than 12 Hz. This suggests that here may exist a subclass of DA cells that exhibits fast firing as proposed in [35] where they the identified ‘a type of dopaminergic neuron within the mesocorticolimbic dopamine system with unconventional fast-firing properties.’ In [35], the observed fast-firing DA neurons were able to sustain higher firing rates above the conventional upper limit of 10 Hz. It would be interesting to have measurements of the differences of the relative strengths of the L-type Ca^2+^ and SK conductances for the fast firing neurons observed by Lammel et al. Moreover, it would also be of interest to examine the neural firing patterns of VTA DA neurons from slice preparations with apamin administered. With increased applied current, would these neurons similarly or reliably enter a fast firing mode?

The framework we have put forth represents a step forward in capturing DA cell dynamics and could be further improved to include more detailed second messenger signaling processes in order to elucidate further differences between the two principal classes of midbrain DA neurons: those of the ventral tegmental area and the substantia nigra pars compacta. However, the GABA studies in this paper, suggest that the SNc DA neurons likely receive a higher level of GABA inhibition due to the observation that SNc neurons display more pacemaker-like firing patterns than VTA DA neurons. The model framework itself could be extended from this conductance-based single compartment model of DA cell activity to include the detailed dynamics of two prevalent nAChR subtypes (*α*4*β*2 and *α*7) and the averaged-activity of the GABA cells (as developed by Graupner and Gutkin [44]). This new framework would take into account a multitude of scales (i.e., linking the VTA mean field activity, detailed conductance-based representations of DA neurons, and the dynamics of the nAChRs) and could yield further insight to the mechanisms underlying burst firing in vivo and aid in elucidating the biophysical manifestations of nicotine addiction.

## Acknowledgements

The authors would like to thank Mark Humphries (The University of Manchester) for his suggestions that contributed to the development of this work. Support for this work was provided by: ANR MNP “Dopanic” (BSG, AMO), ANR-10-LABX-0087 IEC (BSG), ANR-10-IDEX-0001-02 PSL* (BSG), NeRF postdoctoral fellowship (AMO), CNRS (BSG), INSERM (BSG), and an ENP collaborative grant (BSG). Support from the Basic Research Program of the National Research University Higher School of Economics is gratefully acknowledged by BSG.

## Appendix / Tables

**Table.**
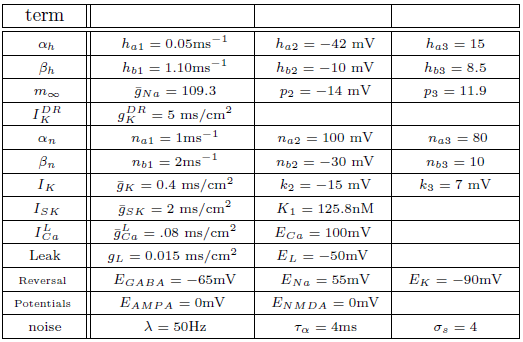
Standard Simulation Parameters

**Table.** Simulation Parameters - II (Depolarization Block)

*Simulation Parameters - II (Depolarization Block)*
| term |  |  |  |
| --- | --- | --- | --- |
| $\beta_h$ | $h_{b1} = 1.5\text{ms}^{-1}$ | $h_{b2} = -11 \text{ mV}$ | $h_{b3} = 13$ |
| $\alpha_n$ | $n_{a1} = 1\text{ms}^{-1}$ | $n_{a2} = 20 \text{ mV}$ | $n_{a3} = 40$ |
| $\beta_n$ | $n_{b1} = 2\text{ms}^{-1}$ | $n_{b2} = -20 \text{ mV}$ | $n_{b3} = 5$ |
| $m_\infty$ | $\bar{g}_{Na} = 109.3$ | $p_2 = -10 \text{ mV}$ | $p_3 = 18$ |
| $I_K^{DR}$ | $\bar{g}_K^{DR} = 18$ | | |
| | $g_{Na}^{pers} = 0.002$ | | $g_L = 0.012$ |

**Table.** Simulation Parameters - III (unveiled)

*Simulation Parameters - III (unveiled)*
| term |  |  |  |
| --- | --- | --- | --- |
| $\alpha_n$ | $n_{a1} = 1\text{ms}^{-1}$ | $n_{a2} = 70 \text{ mV}$ | $n_{a3} = 100$ |
| $\beta_n$ | $n_{b1} = 2\text{ms}^{-1}$ | $n_{b2} = -20 \text{ mV}$ | $n_{b3} = 5$ |
| noise | $\lambda = 25\text{Hz}$ | $\tau_\alpha = 4\text{ms}$ | |

